# Functionally distinct ERAP1 and ERAP2 are a hallmark of HLA-A29-(Birdshot) Uveitis

**DOI:** 10.1101/338228

**Authors:** Jonas J.W. Kuiper, Jessica van Setten, Matthew Devall, Mircea Cretu-Stancu, Sanne Hiddingh, Roel A. Ophoff, Tom O.A.R. Missotten, Mirjam van Velthoven, Anneke I Den Hollander, Carel B Hoyng, Edward James, Emma Reeves, Miguel Cordero-Coma, Alejandro Fonollosa, Alfredo Adán, Javier Martín, Bobby P.C. Koeleman, Joke H. de Boer, Sara L. Pulit, Ana Márquez, Timothy R. D. J. Radstake

**Affiliations:** Department of Ophthalmology, University Medical Center Utrecht, University of Utrecht, Utrecht, the Netherlands; Laboratory of Translational Immunology, University Medical Center Utrecht, University of Utrecht, Utrecht, the Netherlands; Department of cardiology, University Medical Center Utrecht, University of Utrecht, Utrecht, the Netherlands; Department of Genetics, Center for Molecular Medicine, University Medical Center Utrecht, Utrecht University, the Netherlands; Department of Psychiatry, Brain Center Rudolf Magnus, University Medical Center Utrecht, The Netherlands; Department of Human Genetics, David Geffen School of Medicine, University of California, USA; Center for Neurobehavioral Genetics, Semel Institute for Neuroscience and Human Behavior, University of California, USA; The Rotterdam Eye Hospital, Rotterdam, The Netherlands; Department of Ophthalmology, Donders Institute for Brain, Cognition and Behaviour, Radboud university medical centre, Nijmegen, the Netherlands; Department of Human Genetics, Donders Institute for Brain, Cognition and Behaviour, Radboud University Medical Center, Nijmegen, The Netherlands; Centre for Cancer Immunology, Faculty of Medicine, University Hospital Southampton, Southampton, UK; Ophthalmology Department, Hospital de León, IBIOMED, Universidad de León, León, Spain; Ophthalmology Department, BioCruces Health Research Institute, Hospital Universitario Cruces, Barakaldo, Spain; Ophthalmology Department, Hospital Clinic, Barcelona, Spain; Instituto de Parasitología y Biomedicina “López-Neyra”, CSIC, PTS Granada, Granada, Spain; Li Ka Shing Centre for Health Information and Discovery, Big Data Institute, Oxford University, Oxford, UK; Systemic Autoimmune Disease Unit, Hospital Universitario San Cecilio, Instituto de Investigación Biosanitaria de Granada, Granada, Spain; Department of Rheumatology and Clinical Immunology, University Medical Center Utrecht, Utrecht, The Netherlands

## Abstract

Birdshot Uveitis (Birdshot) is a rare eye condition that affects HLA-A29-positive individuals and could be considered a prototypic member of the recently proposed “MHC-I-opathy” family. Genetic studies have pinpointed the ERAP1 and ERAP2 genes as shared associations across MHC-I-opathies, which suggests ERAP dysfunction may be a root cause for MHC-I-opathies. We mapped the ERAP1 and ERAP2 haplotypes in 84 Dutch cases and 890 controls. We identified association at variant rs10044354, which mediated a marked increase in *ERAP2* expression. We also identified and cloned an independently associated ERAP1 haplotype (tagged by rs2287987) present in more than half of the cases; this ERAP1 haplotype is also the primary risk and protective haplotype for other MHC-I-opathies. We show that the risk ERAP1 haplotype conferred significantly altered expression of *ERAP1* isoforms in transcriptomic data (n=360), resulting in lowered protein expression and distinct enzymatic activity. Both the association for rs10044354 (meta-analysis: OR[95% CI]=2.07[1.58-2.71], p=1.24 × 10(−7)) and rs2287987 (OR[95% CI]: =2.01 [1.51-2.67], p=1.41 × 10(−6)) replicated and showed consistent direction of effect in an independent Spanish cohort of 46 cases and 2,103 controls. In both cohorts, the combined rs2287987-rs10044354 haplotype associated with Birdshot more strongly than either SNP alone (meta-analysis: p=3.9 × 10(−9)). Finally, we observed that ERAP2 protein expression is dependent on the ERAP1 background across three European populations (n=3,353). In conclusion, a functionally distinct combination of ERAP1 and ERAP2 are a hallmark of Birdshot and provide rationale for strategies designed to correct ERAP function for treatment of Birdshot and MHC-I-opathies more broadly.

## Introduction

‘*MHC-I-opathy*’ is a relatively new term proposed to unify a group of severe inflammatory diseases, each characterized by strong association with a unique major histocompatibility complex class I (MHC-I) allele and shared immune characteristics [1]. Recent genetic association studies revealed a vast enrichment for polymorphisms in the *endoplasmic reticulum aminopeptidase* (ERAP) genes, ERAP1 and ERAP2, in patients with *ankylosing spondylitis, psoriasis*, or *Behçet’s disease* [2–5]. Importantly, in these conditions, *ERAP* is in epistasis with the risk alleles of MHC-I allele. However, not all patients with ankylosing spondylitis or Behçet’s disease carry a MHC-I risk allele, an observation that has provoked discussions on this unifying concept [6,7] and whether it reflects shared underlying disease etiology.

The mechanism through which ERAP function contributes to MHC-I-opathies is still under intense discussion and the discovery of the underlying mechanism is hampered by several unique characteristics of these specialized enzymes: the peptidases ERAP1 and ERAP2 closely interact with MHC-I by contributing to the generation and destruction of peptides considered for presentation by MHC-I on the cell surface. This key function indicates that aberrant peptide processing by ERAP may be a root cause for MHC-I-opathies [8]. Similar to MHC-I, genetic diversity of *ERAP* genes is conserved by balancing selection [9]. Balancing selection maintains ERAP2 deficiency in one-quarter of the population [9] and drives the polymorphic *ERAP1* gene to encode various protein haplotypes (allotypes) [10], few of which paradoxically confer or reduce risk for developing MHC-I-opathies [8]. To date, ERAP haplotype studies have been limited to the relatively more common MHC-I-opathies [11–12]. In contrast, Birdshot uveitis or *Birdshot chorioretinopathy* (from here on ‘*Birdshot*’) is a severe, extremely rare eye condition (~1-5 cases/500,000) [13,14] that is characterized by enhanced IL-23-IL-17 pathway activation [15–17], and occurs exclusively in HLA-A29-positive individuals [18]. A previous genome-wide association study in Birdshot highlighted strong involvement of the *ERAP* region on chromosome 5q15 [19]. The unique (diagnostic) prerequisite of HLA-A29 [20] and *ERAP* association may classify Birdshot patients within the MHC-I-opathy cluster and, thus, provides an attractive model to define the genetic architecture of this superfamily of conditions [18].

Therefore, we mapped ERAP1 haplotypes and other key variants at 5q15 across a Dutch discovery cohort and validated our findings in a Spanish replication cohort. We interrogated the downstream implications of associated variants using transcriptomic data, molecular cloning and functional assays.

## Results

### Two independent variants at 5q15 associate with Birdshot

To test for genetic variants that associate with Birdshot, we obtained genotype data for 13 key variants at 5q15 in 84 cases and 890 controls from the Netherlands (**Supplementary Table 1**). Using a logistic regression model, adjusted for sex and the top two principle components, we identified a top association at the T allele of rs10044354 in *LNPEP* (odds ratio (OR) = 2.42[95% CI: 1.69-3.45], p = 1.21 × 10^−6^, **Figure 1** and **Supplementary Table 1**). We accounted for the effect of rs10044354 using conditional analysis, and identified the C allele of rs2287987 (encoding valine at amino acid position 349) in *ERAP1* as a secondary (independent) association to Birdshot (OR = 1.99 [95% CI: 1.36-2.90], p = 3.97 × 10^−4^, **Figure 1**). Due to linkage disequilibrium (LD), several other SNPs across the locus also showed strong association, but conditioning on rs2287987 in *ERAP1* removed the remainder of the association signal in the region (**Supplementary Table 1**).

**Figure 1.**
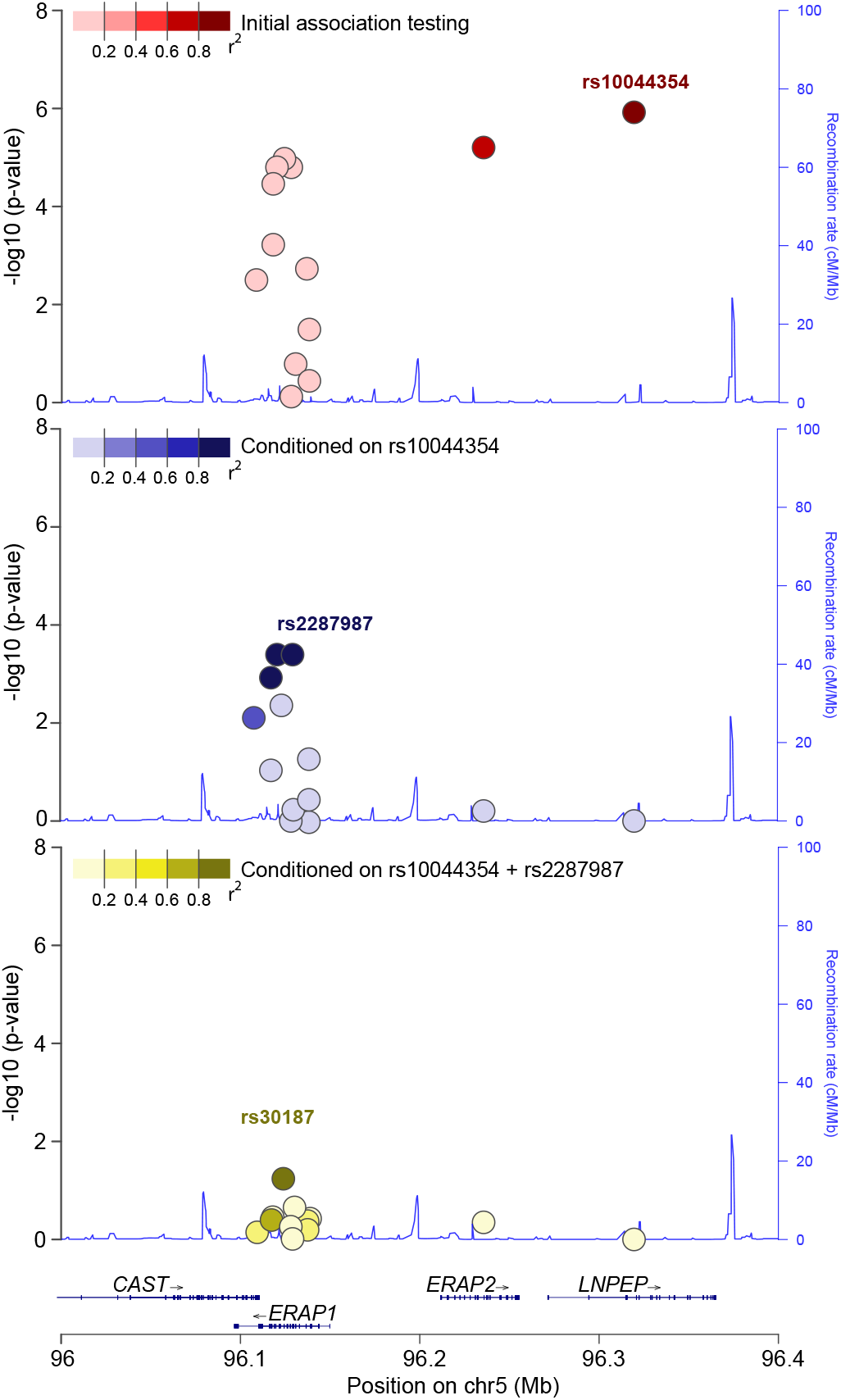
Association and conditional testing for the 13 polymorphisms at 5q15. **A**. Initial association testing mapped the strongest association to rs10044354 in *LNPEP* (Dark red). **B**. Conditioning on rs10044354 revealed independent association for the C allele of rs2287987 (amino acid position 349 in ERAP1) and variants in tight LD (rs10050860 and rs17482075 in dark blue) in *ERAP1*. **C**. Conditioning on both rs10044354 and rs2287987 removed the bulk of the signal (rs30187, p = 0.05). Plot framework was generated by *LocusZoom* [53] and edited for graphical purposes.

We sought to replicate the associations with rs2287987 and rs10044354 using a set of independent samples drawn from an independent Spanish cohort of 46 cases and 2,103 controls. Both associations replicated (**Table 1** and **Supplementary Table 1**) and show consistent direction of effect (meta-analysis, rs10044354, OR [95% CI: =2.07[1.58-2.71], p= 1.24 x 10^−7^; meta-analysis rs2287987, OR [95% CI]: =2.01[1.51-2.67], p = 1.41 x 10^−6^). Interestingly, the combined haplotype rs2287987-/rs10044354 showed disease association exceeding the association for each individual SNP in both cohorts (**Supplementary Table 2**). The strongest association mapped to the protective haplotype rs2287987-T/rs10044354-C (meta-analysis, OR [95% CI]: =0.39 [0.30-0.55],p=3.9 x 10^−9^), closely followed by the risk haplotype rs2287987-C/rs10044354-T (OR [95% CI]: =2.75 [1.92-3.92], p= 2.6 x 10^−8^), the latter present in 50% of all 130 cases and 26% of all 2,993 controls. The frequency of the CT-risk haplotype (0.24 in Dutch cases and 0.19 of Spanish cases) is ~5-6x less common in African (0.027) and Asian (0.021) populations compared to the European population (0.13, **Figure 2A**).

**Figure 2.**
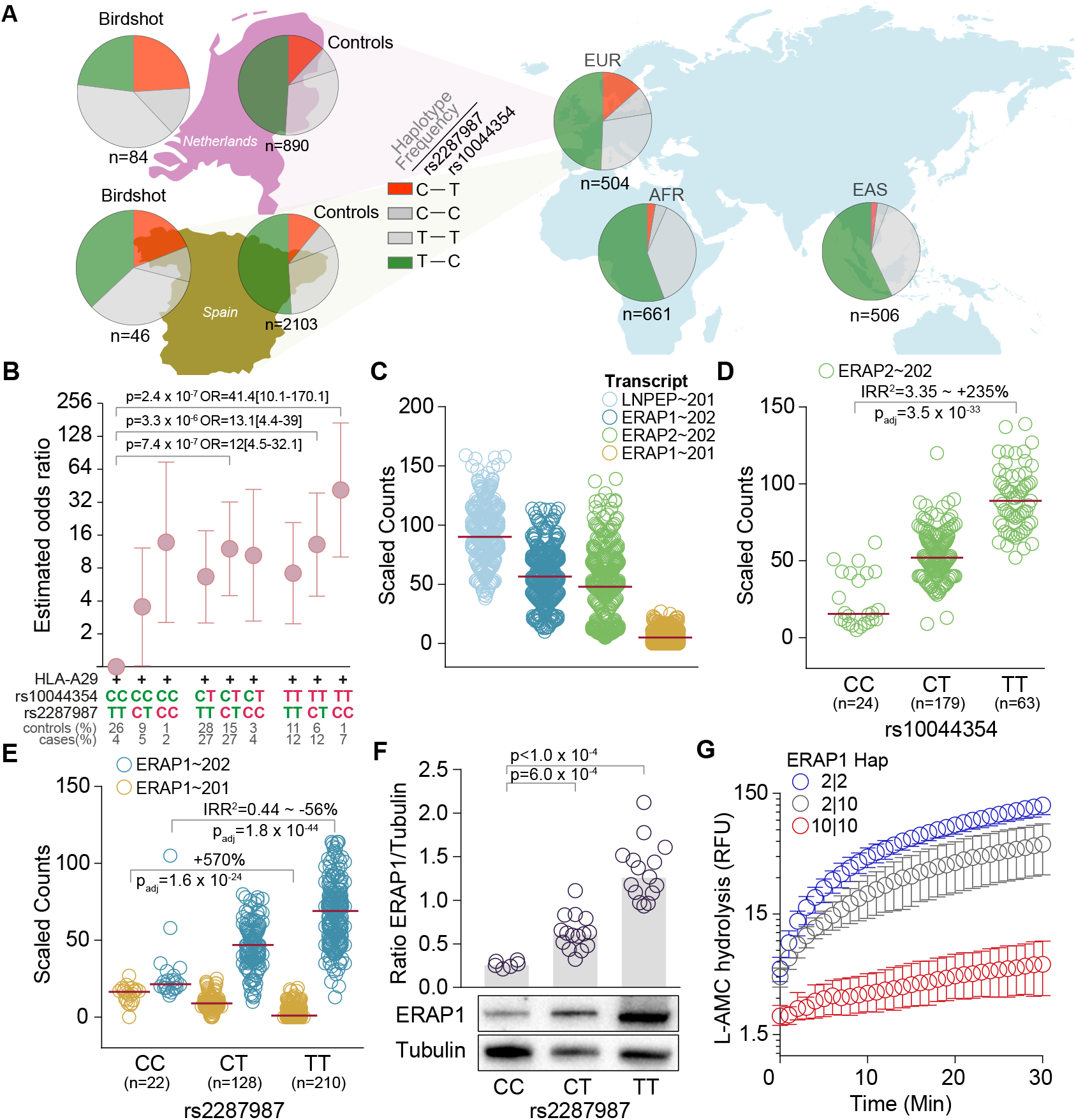
Birdshot associated variants affects ERAP1 and ERAP2 expression and function. **A**. The frequency of the rs2287987-rs10044354 haplotypes in Dutch and Spanish cases and controls, and the EUR, EAS, and AFR super populations of the 1000 Genomes project [48]. The frequency of the haplotypes is indicated as the percentage of all haplotypes for each population. **B**. We tested for association of rs2287987-rs10044354 genotypes in 129 HLA-A29-positive cases and 439 HLA-A29-positive controls in the combined Dutch and Spanish samples. Odds ratios (and 95% confidence intervals) for each genotype combination were calculated relative to lowest risk genotype: the rs2287987-TT/rs10044354-CC (set to an odds ratio of 1). The risk allele (red) and protective alleles (green) are highlighted. **C**. Expression data for transcripts at 5q15 in 360 lymphoid cell lines from the European populations from the GEUVIDAS consortium [48]. Red lines indicate the median expression **D**. The lead SNP rs10044354 is a strong eQTL for ERAP2 independent of rs2248374 (details eQTL analysis in **Supplementary Table 4**). **E**. The Birdshot-associated rs2287987 is associated with changes in the two major ERAP1 transcripts. F. Quantitative Western blot analysis of the protein expression of ERAP1 in HLA-A29-positive cell lines according to the distribution of rs2287987 (more details see **Supplementary Figure 4**). **G**. Hydrolysis (expressed as relative fluorescent units [RFU] of the substrate *Leu-AMC* by immuno-precipitated ERAP1 protein from lymphoid cell lines homozygous *Hap10* (risk haplotype), homozygous *Hap2* (protective haplotype) and a heterozygous donor for these two *ERAP1* haplotypes. Error bars indicate the mean (range) of three independent experiments.

**Table 1:**
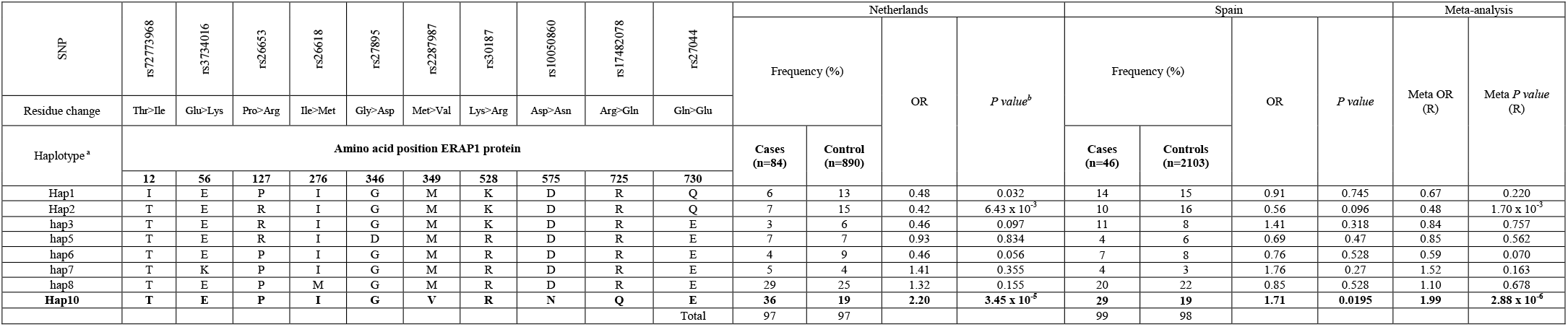
Common ERAP1 haplotypes based on 10 missense variants of 260 Birdshot haplotypes and 5986 control haplotypes from The Netherlands and Spain. ^a^) Haplotype numbers according to *Ombrello et al., 2015* [21]. The 8 haplotypes listed account for >95% of all the haplotypes in the Dutch and Spanish controls. ^b^) Significance (p<1.49 × 10^−4^, see methods). Only Hap10 is significant. We also note a lower frequency of Hap2 in cases.

Since Birdshot is associated with HLA-A29, we investigated the association for the rs2287987-rs10044354 haplotype in the context of HLA-A29 status in the combined 129 HLA-A29-positive cases and 439 HLA-A29-positive controls. Odds ratios for each haplotype’s association to Birdshot were calculated relative to the haplotype carrying the two lowest risk genotypes: rs2287987-TT-rs10044354-CC. In the HLA-A29-postive population, only genotypes that harbor risk alleles from both rs2287987 and rs10044354 associated with Birdshot (p<1.49 × 10^−4^), with the strongest association observed for the rs2287987-CC-rs10044354-TT genotype (OR [95% CI]: = 41.4 [10.08-170.08], p=2.4 x 10^−7^) (**Figure 2B**). Considered collectively, our association testing demonstrates that rs10044354 in *LNPEP* and rs2287987 (i.e., *Hap10*) of *ERAP1* explained the bulk of the association with Birdshot at 5q15 and that combinations of these ERAP variants significantly increase the risk in the HLA-A29-positive population.

### ERAP1 Hap10 is associated with Birdshot

Naturally occurring ERAP1 protein haplotypes have been previously linked to MHC-I-opathies [8]. Considering 10 amino acid variants in *ERAP1* together [21], we identified 8 common (frequency >1%) haplotypes (**Table 1**). The Birdshot-associated C allele of rs2287987 (349V) resides in the ERAP1 haplotype previously labelled ‘*Hap10’* [21]. Indeed, testing for association of phased haplotype data revealed that *Hap10* was associated with Birdshot and replicated in both cohorts (meta-analysis OR = 1.99 [95% CI: 1.49-2.65], p = 2.88 x 10^−6^, **Table 1**). We also noted that *Hap2* was less frequent in cases (**Table 1**). Sixty-three percent of Dutch cases and 46% of Spanish cases had at least one copy of *Hap10* compared to 37.2% and 34.7% of controls, respectively (**Supplementary Table 3**).

### The primary association signal at 5q15 associates with increased ERAP2 expression

Next, we aimed to predict the local transcriptional regulation of the Birdshot-associated loci at 5q15 using RNA-sequencing data from 360 individuals of European ancestry and 82 individuals from African ancestry. We used lymphoid cell lines due to their high expression of ERAPs and representation of expression quantitative trait loci (eQTLs) cis effects of blood cell populations (**Supplementary Figure 6**). We applied a linear mixed model controlling for sex, ancestry, and batch to model if the SNPs imposed eQTLs effects on transcripts encoded by 5q15 (**Figure 2C**); The leading SNP (rs10044354) in *LNPEP* was not associated with the expression of its major transcript (**Supplementary Table 4**). We extracted common missense and splice region variants of *LNPEP* in high LD with rs10044354 from the European populations of the 1000 Genomes [22] and identified the splice region variant rs3836862 (r^2^>0.96 in 5 populations) located near the exons encoding the zinc-binding motif of the enzyme and is predicted to influence splicing. However, we detected no evidence for potential alternative splicing of *LNPEP* by this variant in lymphoid cell lines (**Supplementary Figure 5**). Correcting for the expression of ERAP2 mediated by rs2248374, we found the risk allele of rs10044354 to be strongly associated with *ERAP2* in sample from European Ancestry. Further, this allele associated to a 235% increase in expression of *ERAP2* independent from rs2248374 (IRR^2^ [95% CI]= 3.35[2.76-4.06], p = 3.48 x 10^−33^, **Supplementary Table 4** and **Figure 2D**). In line with this, conditioning on rs2248374 in *ERAP2* in the 130 cases and 2,993 controls combined, we found the association for rs10044354 with Birdshot to be independent from rs2248374 (p = 9.1 x 10^−4^).

### ERAP1 risk alleles reside in a relatively low expressed haplotype

The Birdshot risk variant rs2287987 (349V in ERAP1) is in near complete LD (r^2^=0.99) with rs10050860 (575N in ERAP1, p = 1.46 x 10^−6^ for association with Birdshot). The SNP rs10050860 was recently suggested to represent a splice interfering variant (rs7063), which was linked to changes in the proportion of expression of two major ERAP1 isoforms [23]. We explored the rs7063 variant using rs1057569 (r^2^=1). Indeed, we observed strong association between the rs1057569 genotype and the expression of the two major *ERAP1* transcripts in samples of European ancestry (**Supplementary Table 4**). This variant is also among the top eQTLs for *ERAP1* in lymphoid cell lines investigated in the GTEx project (**Supplementary Figure 7**). The rs2287987 is in moderate LD with rs1057569 (r^2^ = 0.55) and associated with ~50% change in expression of the major ERAP1 isoform (**Supplementary Table 4** and **Figure 2E**), which results in ~5x lower cumulative levels of ERAP1 protein in HLA-A29-positive individuals homozygous for the C allele of rs2287987 (**Figure 2F**). However, the eQTL effects of rs2287987 on *ERAP1* transcripts were completely abrogated once adjusted for rs1057569 (**Supplementary Table 4**). Therefore, we tested whether the rs2287987 disease signal was mediated by rs1057569. Conditioning on rs1057569, we found the rs2287987 signal to be modestly affected (p_meta_ = 4.61 x 10^−4^), while conditioning on rs2287987 neutralized the signal (p_meta_ = 0.39). These results indicate that the association signal at rs2287987 represents a signal beyond the association with the splice interfering variants in *ERAP1*. We note that the A allele of rs1057569 (T allele of rs7063) associated with low expression of ERAP1 protein co-occurred >90% of *Hap10* and *Hap6*, and <10% of the other ERAP1 protein haplotypes in controls. In the Nigergian (YRI) population (n=82), the rs2287987 genotype is not in LD with rs1057569 (r^2^= 0.009), and not linked to ERAP1 expression (transcript ENST00000443439.6, IRR^2^ [95% CI]= 0.93[0.65-1.32], p = 0.68).

### Full-length ERAP1 sequencing of the risk haplotype

We cloned the full-length coding sequence of *ERAP1* from two patients that were heterozygous or homozygous for *Hap10* to reveal the amino acid residues for additional reported amino acid positions in ERAP1 [24]. Analysis of the coding sequence revealed that both copies (based upon 9 individual clones) of *Hap10* encoded the ancestral alleles for additional reported amino acid positions (**Supplementary Table 5**). All detectable transcripts contained a C-terminus identical to the 19-exon transcript ERAP1-202 (ENST00000443439).

### Disease-associated ERAP1 displays distinct enzymatic activity

Considering the association of rs2287987 exceeded variants that govern *ERAP1* expression and that rs2287987 has previously been shown to affect trimming properties of recombinant ERAP1 [25,26], we assessed the relative enzymatic activities of ERAP1 haplotypes *in vitro*. We compared normalized concentrations of ERAP1 protein precipitated from lymphoid cell lines of patients and controls that carry the *Hap10* and *Hap2*, the latter haplotype showing evidence for protection against Birdshot (**Table 1**). Total cellular ERAP1 protein appeared to have slower rates for enzymatic trimming of the archetypical fluorogenic substrate *L-Leucine-4-methylcoumaryl-7-amide* (Leu-AMC) with each sequential increase in Hap10 compared to homozygous carriers of *Hap2* (**Figure 2G**). These results confirm that changes of enzymatic properties in ERAP1 are mediated by haplotypes that carry the C allele of rs2287987 (i.e., *Hap10*), thereby governing functional consequences for patients with Birdshot.

### Non-random distribution of ERAP1-ERAP2 haplotypes in the European population

*ERAP2* encodes two common haplotypes [9]: the A allele of rs2248374 tags Haplotype A [HapA], which encodes a canonical protein, while the G allele tags *HapB*, which lacks functional protein (**Supplementary Figure 8**). Since ERAP1 is postulated to be complemented by ERAP2 in trimming peptides [27], we hypothesized that the expression of ERAP2 protein could be dependent on the *ERAP1* haplotype and that this relationship should be evident in the population. To explore this hypothesis, we used phased genotype data of three populations of 3,353 controls to test whether the frequency of *HapA* of *ERAP2* differed between *ERAP1* haplotypes encoded along the same chromosome. We observed a non-random distribution of *ERAP2-HapA* across the *ERAP1* haplotypes for each European population (**Figure 3** and **Supplementary Table 6**). This reflects the complex pattern of LD (D’) between amino acid variants in *ERAP1* and rs2248374 in *ERAP2* (**Supplementary Figure 9**). A similar non-random distribution was observed for *ERAP1* haplotypes based only on five amino acids [12] (**Supplementary Figure 10**). This demonstrates that the co-occurrence of functional ERAP2 protein is dependent on the ERAP1 haplotype background.

**Figure 3.**
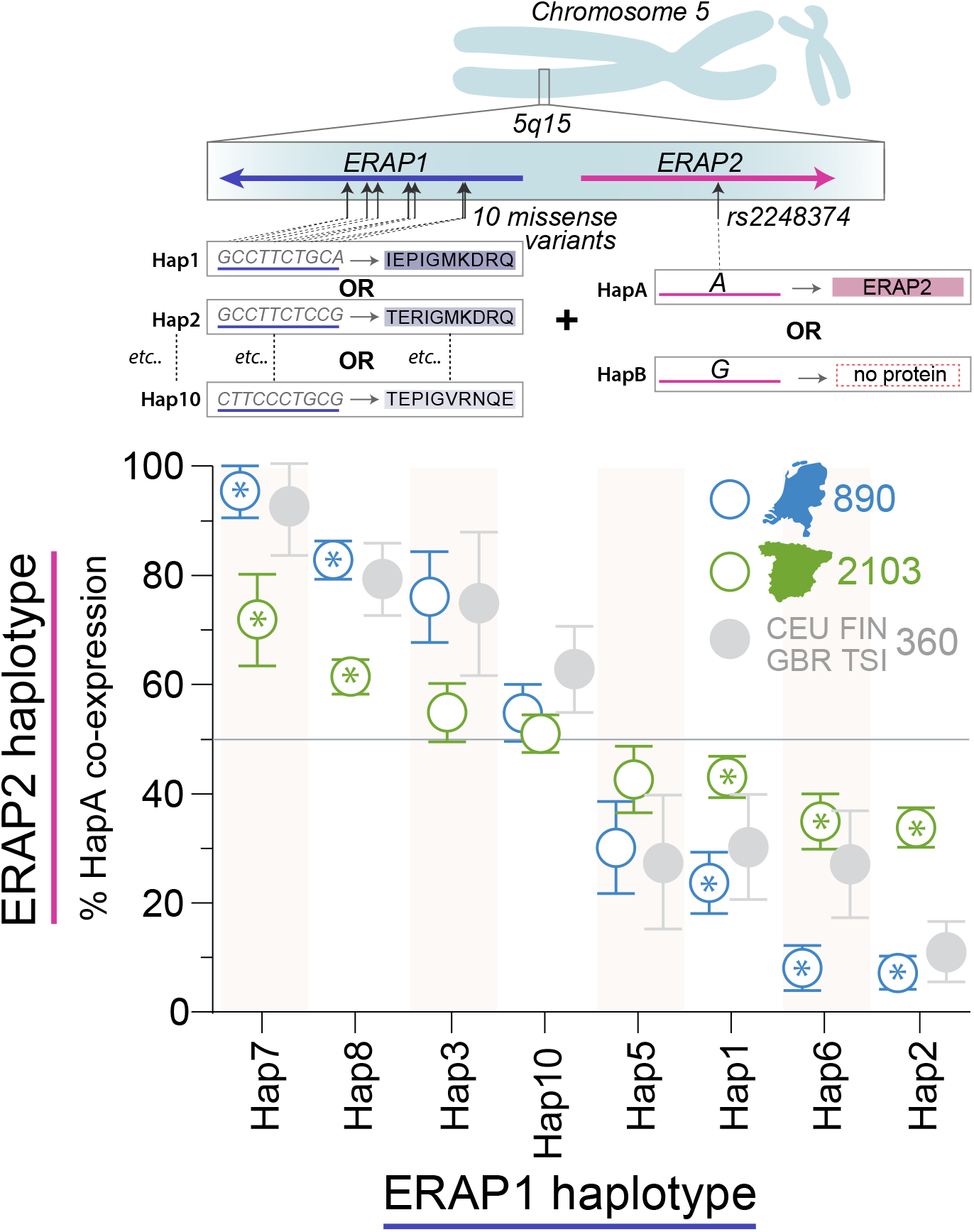
Non-random distribution of the ERAP2-protein coding haplotype across common ERAP1 haplotypes. Ten missense variants in *ERAP1* encode eight discrete and common haplotypes, while *ERAP2* encodes one common protein-coding haplotype (*HapA* tagged by the A allele of rs2248374) and a non-coding haplotypes (tagged by the G allele). Each chromosome 5 harbors one ERAP1 haplotype and one ERAP2 haplotype. Using phased ERAP1-ERAP2 haplotype data of 890 Dutch controls (1,780 phased haplotypes) and 2103 Spanish controls (4,206 phased haplotypes), we outlined the percentage of *HapA* per ERAP1 haplotype on the same chromosome. Error bars indicate the 95% confidence interval based upon 20,000 bootstrap samples. Frequency of *HapA* of ERAP2 for each of the ERAP1 haplotypes was considered non-random if the observed frequency deviated from an expected 50% using an exact binominal test with [A or G rs2248374 for 8 ERAP1 haplotypes = 0.05/16] p<2.5 x 10^−3^ in both the Dutch and Spanish populations with consistent direction of effect. ERAP1 haplotypes that fulfilled these criteria are highlighted by asterisks (for summary statistics see **Supplementary Table 6**). As a reference, we plotted the ERAP1-ERAP2 haplotype data for the 360 samples from the CEU, FIN, GBR, and TSI populations from the 1000 Genomes sample collection used for functional investigation in this study [48].

## Discussion

We defined the common *ERAP1* and *ERAP2* haplotype structure in Birdshot, we confirmed the genetic association with rs10044354 [19] and showed that despite strong LD between rs10044354 and splice region variant rs2248374 in *ERAP2*, the strong eQTL effects of rs10044354 on ERAP2 expression are independent from rs2248374. This finding suggests that a change in expression, and not merely the presence of ERAP2, confers risk for Birdshot and other ERAP2-linked conditions [28]. Thus, future treatment strategies to dampen ERAP2 expression may complement current research efforts focused on the pharmacological inhibition of its aminopeptidase activity [29].

We revealed a second independent association for rs2287987 in *ERAP1*, a SNP that nearly always resides in the common ERAP1 haplotype *Hap10* [21]. The rs2287987 risk genotype is accompanied by changes in *ERAP1* transcript levels and lower protein expression as a result of LD with rs1057569, a strong eQTL for *ERAP1* (**Supplementary Figure 6**). The functional implications of ERAP1 expression need further investigation, especially since the risk genotype linked to *ankylosing spondylitis* (AS) results in increased ERAP1 expression [23]. However, the association with rs2287987 was independent from rs1057569, suggesting disease modifying effects beyond the expression of ERAP1. Indeed, we ascertained that the Birdshot-associated allele of rs2287987 (349V located near the zinc-binding motif of the enzyme) resides in an haplotype that within cells exhibits distinct enzymatic activity and our data support previous studies indicating that Hap10 can be considered functionally antagonistic to Hap2 and Hap1 [24,26]. Also, direct comparison by molecular cloning revealed that *Hap10* in Birdshot and a previously investigated 15-amino acid ERAP1 haplotype, termed \**001*, represent identical ERAP1 haplotypes [10,24]. The *001 (Hap10) was previously shown to provide suboptimal peptide cargo for MHC-I, by reduced trimming of peptides [10,24]. Therefore, Hap10 may confer risk by incomplete destruction of immunogenic epitopes that bind HLA-A29. Eye tissues of Birdshot patients are infiltrated by CD8+ T cells, but to date, evidence for their direct involvement is lacking [18]. ERAP1 and ERAP2 may perhaps also influence T or NK cells function via *Killer immunoglobulin-like* (KIR) receptors, a class of immune receptors that interact with MHC-I that have been genetically linked to Birdshot [30].

Hap10 is also the major risk ERAP1 haplotype for patients with *Behçet’s Disease* (BD) and is highly protective against AS [11,12]. Hap10 is associates with BD assuming a recessive model (Birdshot, p_meta-recessive_ = 9.81 × 10^−5^). Hap1/Hap2 are risk haplotypes for AS and psoriasis [8,12], while being protective for BD and Birdshot. Therefore, Hap10 and Hap2 of ERAP1 play a central role for patients within the MHC-I-opathy cluster: patients with psoriasis have an increased risk of uveitis [31]. Patients with AS and BD often have uveitis [11,42], and the *ERAP1* associations are much stronger in cases that developed uveitis compared with cases with systemic disease alone [2,32]. Although HLA-A29 is also common in some non-European populations, Hap10 and particularly the Hap10-ERAP2 (rs2287987-rs10044354) haplotype are less common in Asian and African individuals compared to European and, as we show, if present may exhibit different functional effects on ERAP1, which may explain why Birdshot primarily affects patients of European descent [18].

Although 90% of all cases carried either a risk allele for rs2287987 or rs10044354, 50% of all cases were positive for the combined haplotype. A proportion of Birdshot patients may have self-limiting disease, while others will develop irreversible retinal dysfunction and blindness [13]. It will be interesting to evaluate the potential relation between *ERAP* haplotypes and clinical outcome (e.g., electroretinography) in follow-up studies. This was not possible because of the relatively limited sample size, owing to low prevalence of this rare disease [13,14]. This limitation also hampered the investigation for association with other SNPs with evidence for more moderate effect size (e.g. rs30187). Regardless, the strongest associations for Birdshot include the rs2287987-rs10044354 haplotype and HLA-A29, a triad of loci that encode functionally tightly related proteins of the class I antigen presentation pathway and warrants further investigation into antigen presentation in human uveitis. Whether ERAP1 and ERAP2 contribute to disease by direct interaction in the endoplasmic reticulum is not known [43–45]. In the absence of ERAP2 protein, cells with distinct *ERAP1* haplotypes show substantial differences in HLA-A29 peptidome composition, which demonstrates that ERAP1 is able to directly influence HLA-A29 [43]. We recently showed that ERAP2 increases the number of 9-mer peptides presented by HLA-A29 [36]. This suggests that some peptides bound by ERAP2 may be protected from destruction by ERAP1, which will be further facilitated in a hypoactive ERAP1 background (i.e., Hap10). These results favor our hypothesis that a limited set of uveitogenic peptides drive T cell mediated pathology in Birdshot [18]. However, ERAP2 has different peptide preferences compared to ERAP1 [27,29] and MHC-I peptidome studies support separate contributions of Hap10 and ERAP2 in manipulating the peptide composition presented on the cell surface [35]. This reflects the lack of LD between the protein-coding *ERAP2* haplotype and MHC-I-opathy linked Hap10 and Hap2/Hap1 in the European populations investigated. However, ERAP2 protein expression more often occurs in *ERAP1* haplotypes Hap8 and Hap7 (and perhaps Hap3) background, as a result of LD (D’) between SNPs including rs3734016, and rs26618 and rs2248374 (**Supplementary Figure 9**). The lack of detailed MHC-I data hampered further investigation of this phenomena, because ERAP2 may not equally contribute to the peptidome of each MHC-I molecule [26,34]. However, this would demand a large population study with sufficient power to investigate the ERAP1-ERAP2 haplotypes for distinct MHC-I haplotypes. Such studies will also be key to advance our understanding of ERAP function in inflammation, cancer, and less explored fields such as transplantation biology and vaccine efficacy studies.

The original concept of the MHC-I-opathy aims at clustering subsets of clinical phenotypes on the basis of one or multiple unifying molecular fingerprints [1]. Here, we explicitly refer to the subgroup of patients that carry risk MHC-I alleles (and ERAP variants in epistasis) when referring to ‘MHC-I-opathy’ patients. Our results in Birdshot further substantiate the existence of such concept. We envision that molecular reclassification based on a patient’s MHC-I and ERAP haplotype may improve clinical decision making as well as drug discovery (e.g. ERAP inhibitors) in the near future. Emerging studies explore this concept, those of which that have linked clinical response to *ERAP* variants [37–40]. Ultimately, understanding how to pharmacologically influence ERAP function [41] may lay the foundation for the design of effective therapeutic strategies to correct ERAP function in accordance with the patient’s molecular fingerprint to tailor the treatment of MHC-I-opathy patients specifically.

## Methods

### 5q15 variant and haplotype testing in Dutch and Spanish cohorts

This study was performed in compliance with the guidelines of the Declaration of Helsinki and has the approval of the local Institutional Review Board (University Medical Center Utrecht). Genotype data for 13 key variants at 5q15 (**Supplementary Table 1**) and HLA-A29 status from 84 Dutch cases and 890 Dutch controls from European ancestry were obtained from a previously-described discovery collection and the Genome of the Netherlands Project (GoNL) release 4 [19,42]. The number of samples is slightly fewer than the Dutch discovery cohort because some samples no longer met the criteria for Birdshot Uveitis (e.g., presence of systemic disease) or failed to pass quality control requirements. Imputation of untyped SNPs was conducted using the GoNL and 1000 Genomes Project Phase 3 data (imputed SNPs with info>0.95) [43]. Disease association testing was performed with PLINK v1.07 [44] using a logistic regression model correcting for the top two principal components and sex (**Supplementary Figure 1**). For replication, the genotype data (n=13) (**Supplementary Table 1**) and HLA-A29 status were extracted from 46 Spanish cases and 2,103 Spanish controls from a previously-described non-infectious uveitis collection [45] and tested with a logistic regression model (replication was defined as Bonferroni-corrected p-value for 2 LD blocks at 5q15 = 0.025). Meta-analysis of Dutch and Spanish cohorts was performed using inverse variance-weighted metaanalysis [46]. Common ERAP1 haplotypes (10-amino acid haplotypes, which occur in >1% of controls, see **Table 1** and **Supplementary Figure 2**) were phased using PLINK. The estimated haplotype frequencies for the identified 8 common ERAP1 haplotypes based matched the frequency of haplotypes estimates of genotype data by *SHAPEIT2* (considered the most accurate method available [47]). Therefore, the phasing accuracy of PLINK was considered sufficient for *ERAP1* haplotype association testing using logistics regression (**Supplementary Figure 3**). The total testing burden for SNP association was estimated at ([13 variants + 8 ERAP1 protein haplotypes] x 3 models + 4 rs2287987-rs10044354 haplotypes = 67 independent tests) a 1% type I error rate with p<1.49 × 10^−4^, which was used as a threshold to prioritize SNP associations calculated in the discovery and meta-analysis. We tested for the rs2287987-rs10044354 haplotype in the 129 HLA-A29-positive Dutch and Spanish cases and 439 HLA-A29 positive Dutch and Spanish controls using a logistic regression model in the R software package. The odds ratio for each possible genotype was calculated by exponentiating the coefficient from the logistic regression. The low risk rs10044354-CC/rs2287987-TT genotype was set as the baseline (OR=1). P-values from the logistic regression below p<1.49 × 10^−4^ were considered significant.

### Investigation of the distribution of ERAP1 and ERAP2 haplotypes in European populations

The major haplotype of *ERAP2* is under balancing selection, where the G allele rs2248374 tags a non-protein coding haplotype [HapB] and the A allele of rs2248374 tags the protein-coding haplotype [HapA] [9]. The frequency of the A allele of rs2248374 co-occurring on the same chromosome with each of the eight common ERAP1 haplotypes was determined in the 890 Dutch and 2103 Spanish controls using phased haplotype data using SHAPEIT2 [47], and phased haplotype data were obtained for the British [GBR], Finnish [FIN], Utah residents with Northern and Western European Ancestry [CEU], and Tuscany [TSI] populations from the GEUVIDAS consortium [48]. The 95% confidence intervals of the observed frequencies were estimated using bootstrap sampling (n=20,000). We used a two-sided exact binominal test to assess whether the observed frequencies were non-random (deviation from a test probability of 0.5 or the hypothesis that ERAP2 co-occurs in 50% of any particular ERAP1 haplotype). Frequency distribution of ERAP2 for each of the ERAP1 haplotypes was considered non-random at [A or G rs2248374 for 8 ERAP1 haplotypes = 0.05/16] p<2.5 x 10^−3^ in both the Dutch and Spanish populations (with consistent direction of effect).

### Statistical framework for 5q15 eQTL analysis

To investigate the transcriptional regulation of the variants at 5q15, RNA-sequencing data from the GEUVADIS consortium of 360 lymphoid cell lines of European ancestry and 82 African samples [48] were extracted via the *recount2* package, a resource for expression data for uniformly processed and quantified RNA-sequencing reads from >70,000 human samples [49]. Transcript-level counts of all samples of the GEUVADIS study were scaled using the scale counts function in the recount2 package to take into account differing coverage between samples. Data corresponding to reported transcripts from *ERAP1, ERAP2*, and *LNPEP* (n=25 using *Ensembl* [22] transcript IDs) were extracted for all samples, of which 4 transcripts were sufficiently expressed and considered for testing the effect of genotype on transcript expression (ERAP1~201, ENST00000296754.7; ERAP1~202, ENST00000443439.6; ERAP2~202, ENST00000437043.7; LNPEP~201, ENST00000231368.9). The phased genotype data were obtained via the *Ensembl* portal [22]. To interrogate the effect of genotype on local transcript expression, we fitted a generalized linear mixed model (GLMM) using a negative binominal fit of the count data and logarithmic link using the *lme4* package in R (v3.3.2)[50], with a fixed effect of patients sex and random effects of laboratory (n=7) and population (n=5) of the 1000 Genomes project (formula 1). For pairwise conditioning, the genotype of the studied SNP was added as a covariate into the GLMM (formula 2):

~~~
(1) : glmer.nb(expression ~ genotype + sex +(1|Laboratory)+(1|Population))
(2) : glmer.nb(expression ~ genotype + sex + conditional SNP genotype + (1|Laboratory)+(1|Population))
~~~

Genotypes were ranked using their imputed dosage value (a number from 0 to 2, indicating the number of risk alleles for each SNP; **Supplementary Table 1**) adapted from a previous method [23]. Patients homozygous for the G allele of rs2248374 (n=94) were removed to fit a model for ERAP2 expression in HapA-positive individuals. The incident rate ratio (IRR) is the exponential of the coefficient returned from the GLMM. We used IRR^2^ as the metric for change in transcript expression, which is conceptually similar and close to fold the change in expression (mean count of the transcript) between individuals homozygous for the protective over individuals homozygous for the risk alleles. IRR^2^ <0.60 or >1.30 (>30% change in expression) were prioritized for reporting as biologically relevant. Effect size metrics of eQTL analysis from lymphoid cell lines and whole blood from the *Genotype-Tissue Expression* (GTEx) project Phase 2 for genes at 5q15 were obtained via the GTEx portal [51].

### Western blot analysis of ERAP1 and ERAP2

Human lymphoid cell lines were generated from peripheral blood of Dutch cases recruited at the Departments of Ophthalmology at the University Medical Center Utrecht and Radboud University Medical Center, the Netherlands. Cells were grown in Roswell Park Memorial Institute (RPMI) 1640 medium supplemented with 10% heat-inactivated fetal bovine serum and antibiotics. The following HLA-A29-positive cell lines were obtained from the *National Institute of General Medical Sciences* (NIGMS) Human Genetic Cell Repository at the Coriell Institute for Medical Research: GM19310, GM19397, GM19452, HG00096, HG00113, HG00116, HG01082. Protein lysates (10μg/lane) of >2 × 10^6^ cells were separated on a 4-20% Mini-PROTEAN TGX gel (Bio-Rad Laboratories) and transferred to a Polyvinylidene difluoride (PVDF) membrane and detected using antibodies to ERAP1 and ERAP2 (AF3830, AF2334, both R&D Systems), and α-tubulin (T6199, Sigma), and secondary antibodies conjugated to Horseradish Peroxidase. The ratio of the intensity of the ERAP1 over α-tubulin band was calculated using Image Lab 5.1 (Bio-Rad Laboratories) and grouped according to the distribution of rs2287987 genotype. Details on the Western blot analyses of lymphoid cell lines are outlined under **Supplementary Figure 4**.

### LNPEP sequencing

We sequenced exon 5-8 of *LNPEP* from total RNA (cDNA) in steady-state or 24 hour stimulated (TLR7/8 [R848] and TLR9 [CpG-B ODN 2006] agonists) lymphoid cell lines of 4 individuals homozygous for the deletion and 4 homozygous for the C allele of rs3836862, as determined by sanger sequencing. Detailed methods and used primers are provided in **Supplementary Figure 5**.

### Full-length ERAP1 cloning and sequencing

RNA was isolated from >2 × 10^6^ cells of two patients with Quick-RNA™ MiniPrep kits (Zymo Research) and immediately used to generate cDNA with the Transcriptor High Fidelity cDNA synthesis kit (Roche) using the provided random hexamere primers. *ERAP1* was directly amplified from cDNA as described [10]. The product was gel-purified, and cloned into the blunt-end cloning vector, pCR-Blunt II, using the Zero Blunt ^®^ TOPO ^®^ PCR Cloning kit (Life Technologies) or cloned into StarGate^®^ pESG-IBA vectors (IBA Lifesciences), and sequenced.

### Cellular ERAP1 enzymatic activity

Total cellular ERAP1 was immuno-precipitated with anti-ERAP1 (5ug/mL, AF2334, R&D Systems) antibody and protein G-sepharose beads as described [52]. Equivalent concentrations of protein samples (normalized by Bradford assay) of purified ERAP1 were incubated with 50μM Leucine-aminomethylcoumarin (L-AMC) and fluorescent intensity was followed for 30 minutes with excitation at 365/380nM and emission at 440/450nm.

## Acknowledgement

We would like to thank prof. Aniki Rothova for support in evaluation of Birdshot patients. JJWK and SLP are supported by a VENI Award from the Netherlands Organization for Scientific Research (N.W.O. project numbers 016.186.006, and 016.186.071). AM is recipient of a Miguel Servet fellowship (CP17/00008) from the Spanish Ministry of Economy, Industry and Competitiveness. JJWK was supported by unrestricted grants of the Fischer Stichting, the Landelijke Stichting voor Blinden en Slechtzienden, and the Algemene Nederlandse Vereniging ter Voorkoming van Blindheid, that contributed through *UitZicht*. The funding organizations had no role in the design or conduct of this research. The authors have no conflict of interest to declare.

## Conflict of Interest Statement

The authors state that they have no conflict of interest to disclose.

## Supplementary Figures

**Supplementary Figure 1.**
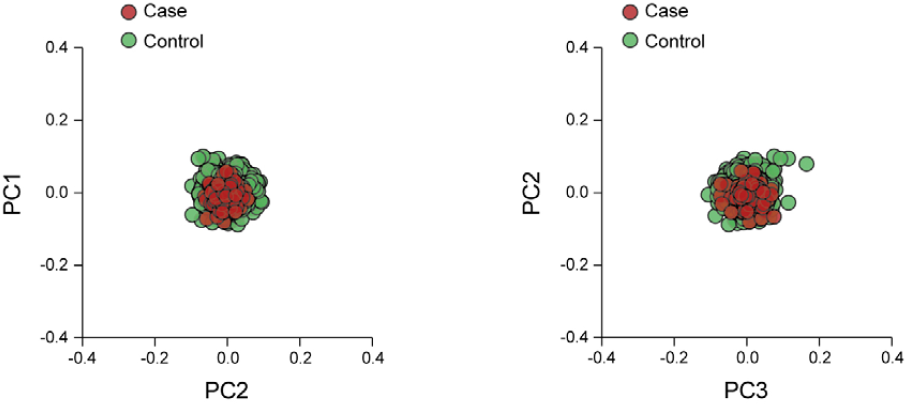
Analysis of genetic matching of cases and controls using principal component analysis (PCA). PCA of the first versus second and second versus third component in cases and controls using genotype data from the Dutch collections as described in the Method section. We filtered SNPs with minor allele frequency (MAF) <0.1, excluding the *MHC*, two large inversions on chromosomes 8 and 17 and the *LCT* locus on chromosome 2. We further filtered out SNPs that are not in Hardy-Weinberg equilibrium (p<10^−4^), or showed a pairwise LD of r^2^ 0.2 or more. PCA was computed using a set of 75,239 random and independent SNPs autosome-wide (chr 1-22) in EIGENSOFT (*Price et al. Principal components analysis corrects for stratification in genome-wide association studies. Nat Genet. 2006 Aug;38(8):904-9*).

**Supplementary Figure 2.**
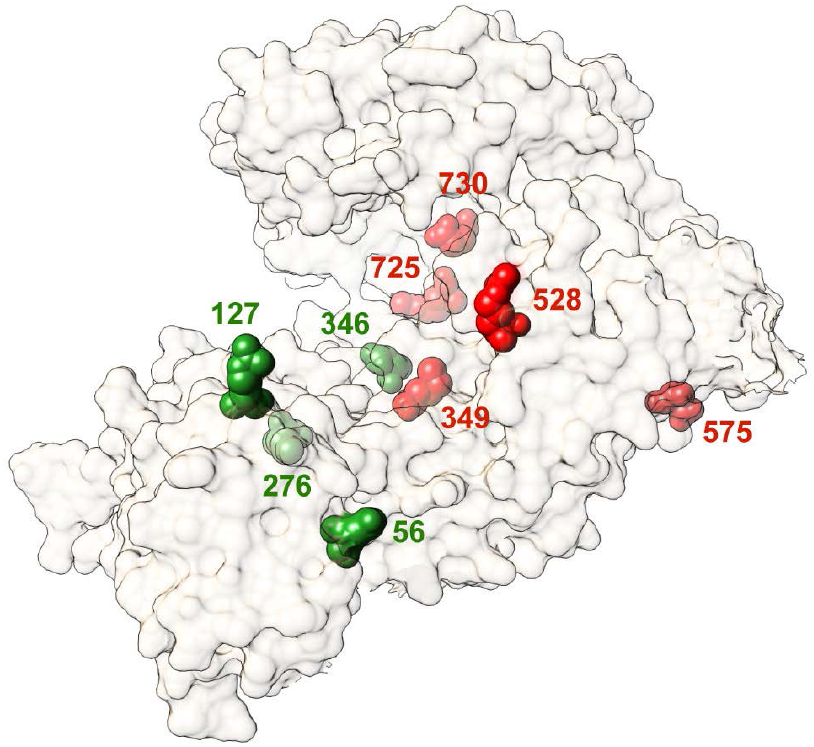
Three-dimensional surface model for the ERAP1 allotype associated with Birdshot uveitis (Protein Data Bank entry: 3MDJ). Common variant amino acid residues in ERAP1 are displayed as spheres; Ancestral amino acids are highlighted in green and non-ancestral amino acid positions are highlighted in red. 3D structure was produced using UCSF Chimera (*Pettersen, E. F. et al. UCSF Chimera - a visualization system for exploratory research and analysis. J. Comput. Chem. 25, 1605–1612 2004*).

**Supplementary Figure 3.**
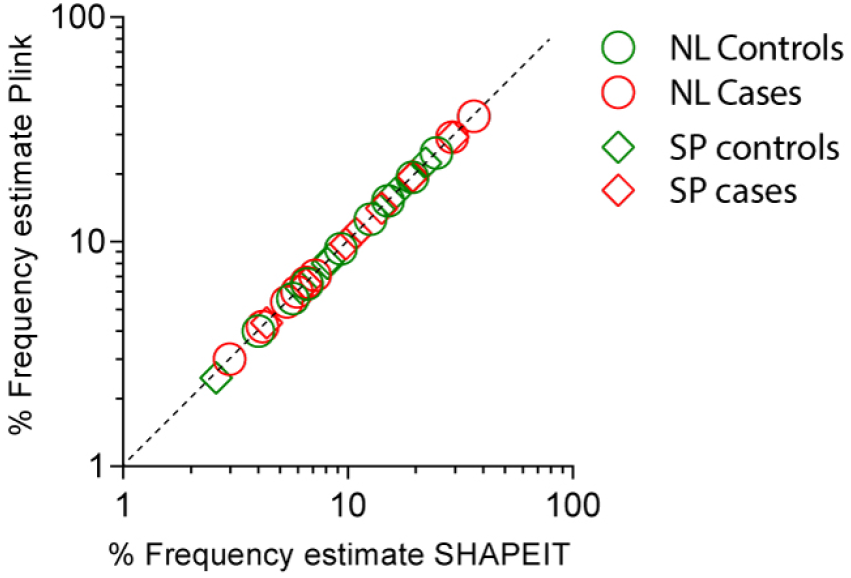
The estimated haplotype frequency (%) for the 8 common ERAP1 haplotypes using Plink [44] and SHAPEIT [47] for cases (in red) and controls (in green). The dotted diagonal line indicates perfect correlation between the estimated haplotype frequencies.

**Supplementary Figure 4.**
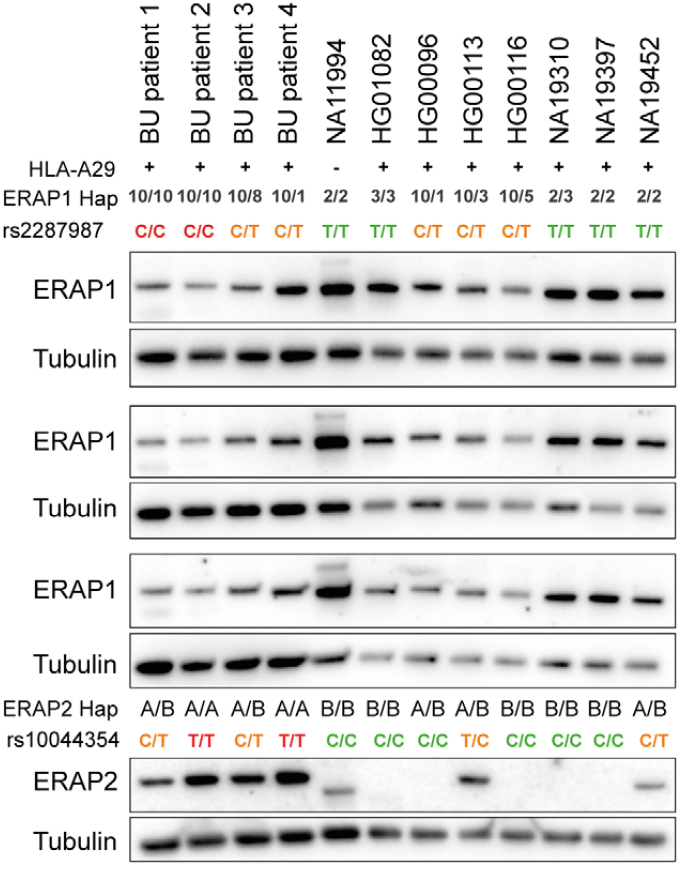
ERAP1 and ERAP2 expression in B cell lines of Birdshot uveitis and HLA-A29-positive controls. We assessed ERAP1 and ERAP2 protein levels in EBV transformed B cells (EBV-LCL), from twelve HLA-A*2902-positive individuals (except NA11994) from the 1000 Genomes project and EBV-LCLs generated from peripheral blood mononuclear cells of four BU patients genotyped in this study, by western blotting. Protein lysates (10μg/lane) from cells were separated on a 4-20% Mini-PROTEAN TGX gel (Bio-Rad Laboratories) and transferred to a PVDF membrane. Proteins were detected using a 1:5000 dilution of primary antibodies; goat anti-ERAP2 polyclonal antibody (AF3830, R&D Systems), goat anti-ERAP1 polyclonal antibody (AF2334, R&D Systems) and anti-α-tubulin monoclonal prepared in mouse (T6199, Sigma). Anti-mouse secondary antibody conjugated to Horseradish Peroxidase (HRP) (Jackson ImmunoResearch; 1:5000) and anti-goat secondary antibody conjugated to HRP (DAKO; 1:5000) were used to probe primary antibodies. Protein bands were detected with Amersham Prima Western Blotting (RPN22361, GE Healthcare) on the ChemiDoc Gel Imaging System (Bio-Rad Laboratories). Independent experiments are individually outlined below for ERAP1. The ratio of the intensity of the ERAP1 over α-tubulin band was calculated using Image Lab 5.1 (Bio-Rad Laboratories) for each experiment. All ratios of the intensities were grouped according to the distribution of *Hap10* (red = homozygous for rs2287987-C, orange = heterozygous for rs2287987-C, green is noncarrier) for quantitative analysis as outlined in **Figure 2F**.

**Supplementary Figure 5.**
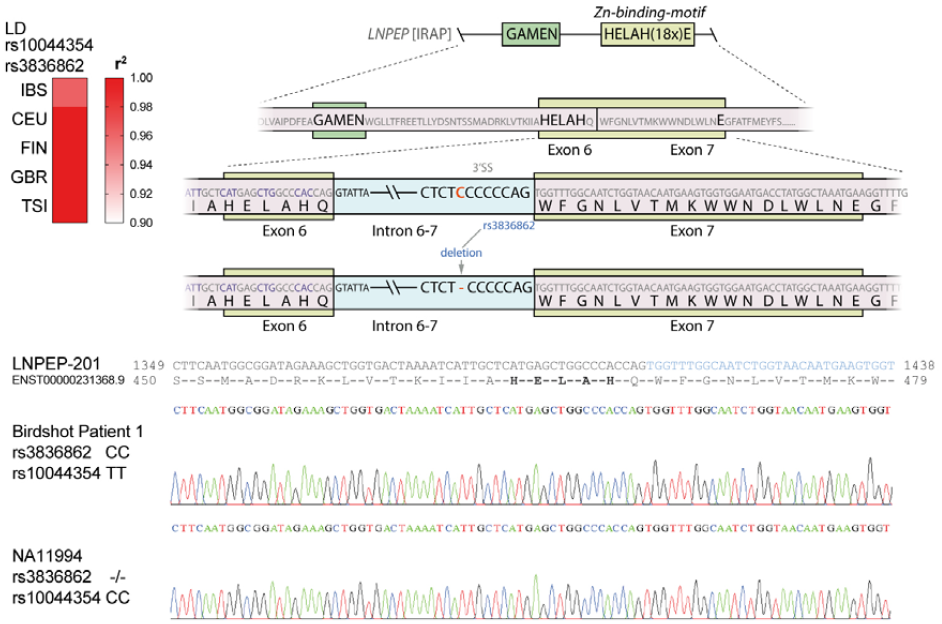
rs10044354 is in tight LD with a splice region variant in *LNPEP*. We searched the 5 European populations (British [GBR], Iberian [IBS], Finnish [FIN], Utah residents with Northern and Western European Ancestry [CEU], and Tuscany [TSI]) of the *1000 Genomes Project* [43] for missense or splice region variants of *LNPEP* in high LD (r^2^ >0.75) with rs10044354; The rs10044354 variant is in tight LD (r^2^ >0.96-1.0) with rs3836862 located in the splice region of intron 6-7 of *LNPEP*. These exons encode the key Zn-binding motif of the enzyme. The *Human splicing finder* software (http://www.umd.be/HSF3/) predicted a potential impact of the rs3836862 on splicing of this region. To explore if rs3836862 altered splicing of intron-6-7, we tested if the presence of rs10044354/rs3836862 affects the sequence of the predominant mature transcripts near this splice region. We selected four lymphoid cell lines that were homozygous null and four homozygous carries of the C allele of rs3836862 (as determined by Sanger sequencing). For each cell line, a total of 1 × 10^6^ cells were cultured in 2 mL RPMI 1640 (Life Technologies), supplemented with 10% (v/v) heat-inactivated FCS (Biowest) and 1% (v/v) penicillin and streptomycin (Life Technologies) in a six wells plate for 24 hours and stimulated with TLR agonists resiquimod (*R848*; 1 μg/ml), or CpG-B (*ODN684*; 5 μM, both from *InvivoGen*), or left untreated. Total RNA was extracted using QiaCube and RNeasy mini kit (*Qiagen*) and a 443 bp cDNA fragment from exon 5 to exon 8 of *LNPEP* was generated by SuperScript ^®^ cDNA synthesis kit (Invitrogen, UK) using 5’-3’ (forward) TATGCCTTGGAAACAACTGTGA and 5’-3’(reverse) TGTTCTGAAGACTGAACAGA TGAT. A 438 bp cDNA fragment was amplified using proof reading Pfu DNA polymerase-PCR reaction (Agilent Technologies, Santa Clara, CA, USA) with 5’-3’(forward) TGCCTTGGAAA CAACTGTGAAG and 5’-3’ (reverse) TCTGAAGACTGAACAGATGATGA. A single band was detected in all conditions by agarose gel and sequenced at Macrogen Europe (Netherland) with 5’-3’(forward) GGAAACAACTGTGAAGCTTCTTGA and 5’-3’(reverse) CTTTCTTCATGGTTTT AAATCGAGC. Chromatograms from representative donors (unstimulated) from each group are indicated below. These analyses detected a sequence identical to the canonical transcript of *LNPEP* (Ensemble ID *ENST00000231368*) in all samples. We concluded that rs3836862 does not affect the sequence of the predominant (detectable) mature LNPEP transcripts in lymphoid cell lines under the described conditions.

**Supplementary Figure 6.**
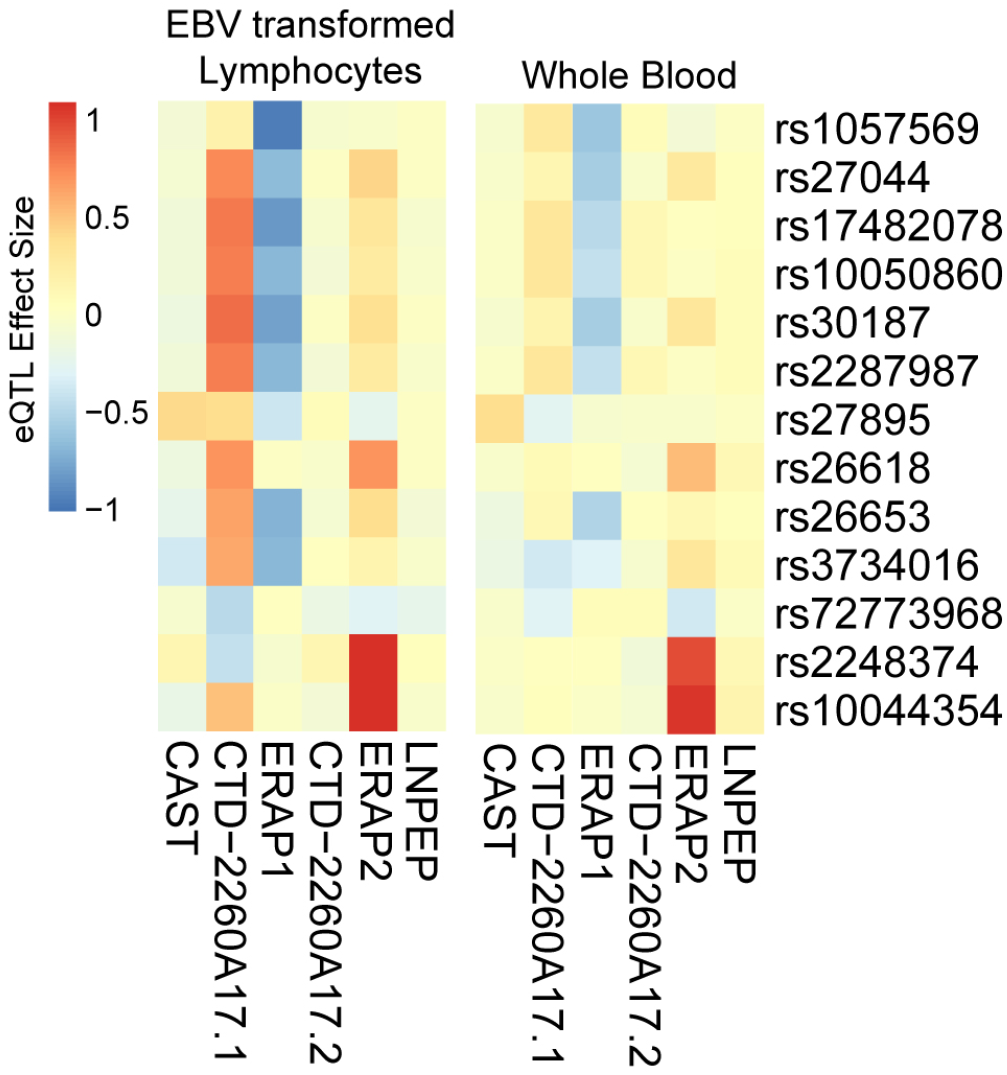
The normalized effect size from the eQTL analysis for variants at 5q15 derived from the Genotype-Tissue Expression (GTEx) Project. Data for 117 lymphoid cell lines and 369 whole blood samples were obtained from the GTExportal (GTEx Analysis Release V7, reference 51) as described in the method section.

**Supplementary Figure 7.**
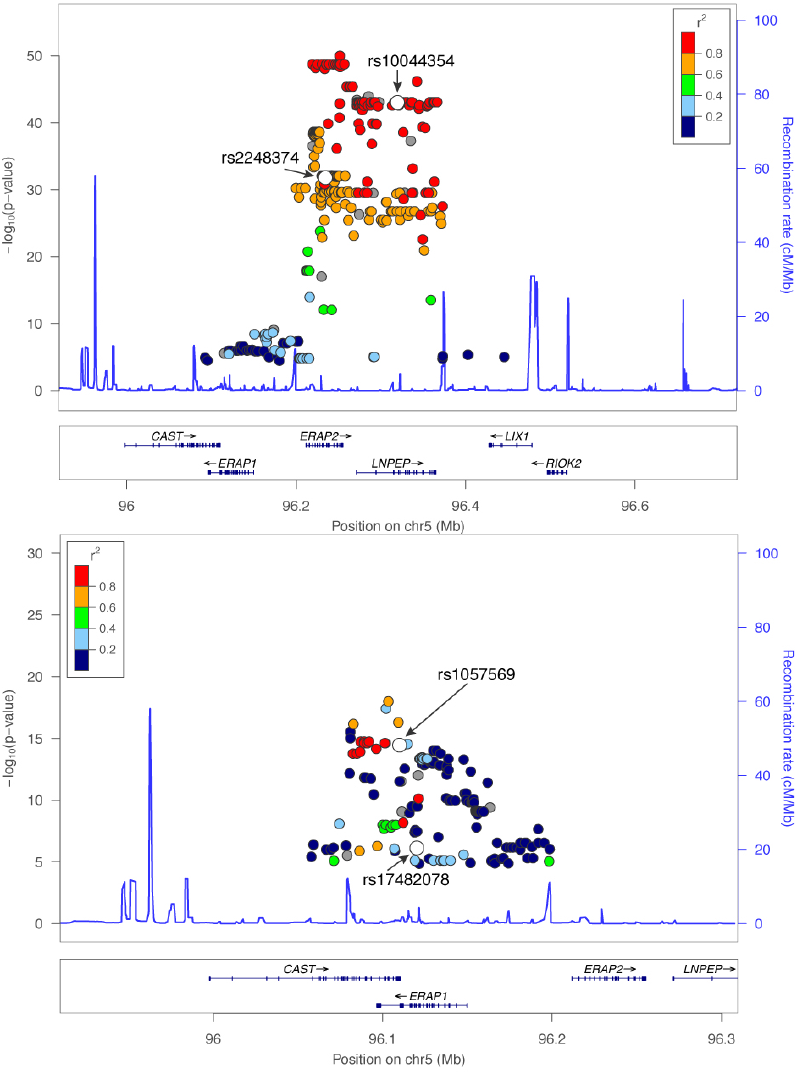
eQTL data for variants in *ERAP1* and *ERAP2* in lymphoid cell lines of 117 individuals from the Genotype-Tissue Expression (GTEx) Project. P-values from eQTL analysis for *ERAP1* and *ERAP2* were obtained from the Genotype-Tissue Expression (GTEx) [51] Project (GTEx Analysis Release V7). Gene expression for *ERAP1* is based on ERAP1~202 (EnSt00000443439, average TPM unit of transcript expression = 29.2) in EBV-LCLs and ERAP1~201 (ENST00000296754 average TPM = 2.16, other transcript TPM<2). Gene expression for ERAP2 based on ERAP2~202 (ENST00000437043, average TPM = 18.1) in EBV-LCLs (other transcripts with >18 exons show TPM<2). The eQTLs rs1057569 for *ERAP1* and rs10044354 for *ERAP2*, and rs2248374 and rs17482078 are indicated in white. The hg19 European [EUR] 2014 1000 Genomes LD structure (r^2^) is color coded. Plots were generated in LocusZoom [53].

**Supplementary Figure 8.**
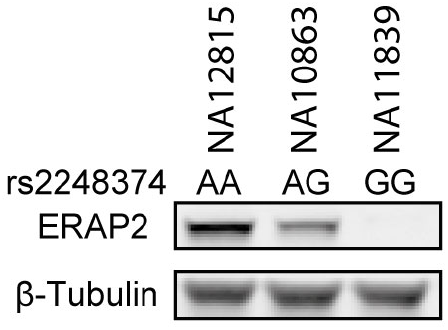
The common splice region variant of *ERAP2* tags the bimodal distribution of protein expression. *ERAP2* encodes at least two common haplotypes that are tagged by rs2248374. The A allele tags the protein-coding haplotype (HapA), while the G allele tags a haplotype that encodes a transcript that is usually prone to none-sense mediated decay [9] and loss strongly reduced protein expression. Expression of ERAP2 in CEPH controls according to rs2248374 genotype is provided for three B-cell lines from individuals from the CEPH panel (*Dausset J. et al. Centre d’etude du polymorphisme humain (CEPH): collaborative genetic mapping of the human genome. Genomics. 1990;6:575-577*). Genotype information was obtained from the *Ensembl Genome Browser* (rs2248374 or rs2549797 that is in full LD [r^2^ =1] with rs2248374). Cells were lysed and ERAP2 expression was assessed after SDS-PAGE and western blotting with antibodies to ERAP2. The endogenous levels of β-tubulin protein were analyzed as a loading control.

**Supplementary Figure 9.**
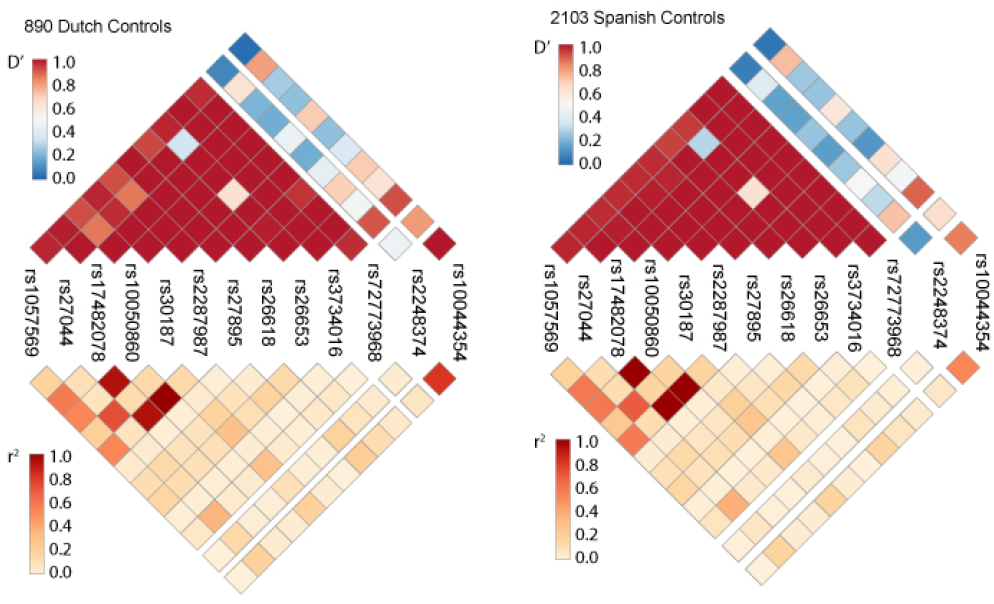
Linkage disequilibrium across the 13 polymorphisms in *ERAP1* and *ERAP2* in cases and controls. The linkage disequilibrium across the functional variants at 5q15. The heatmaps show the D’prime and r^2^ values calculated for SNPs in the Dutch and Spanish controls.

**Supplementary Figure 10.**
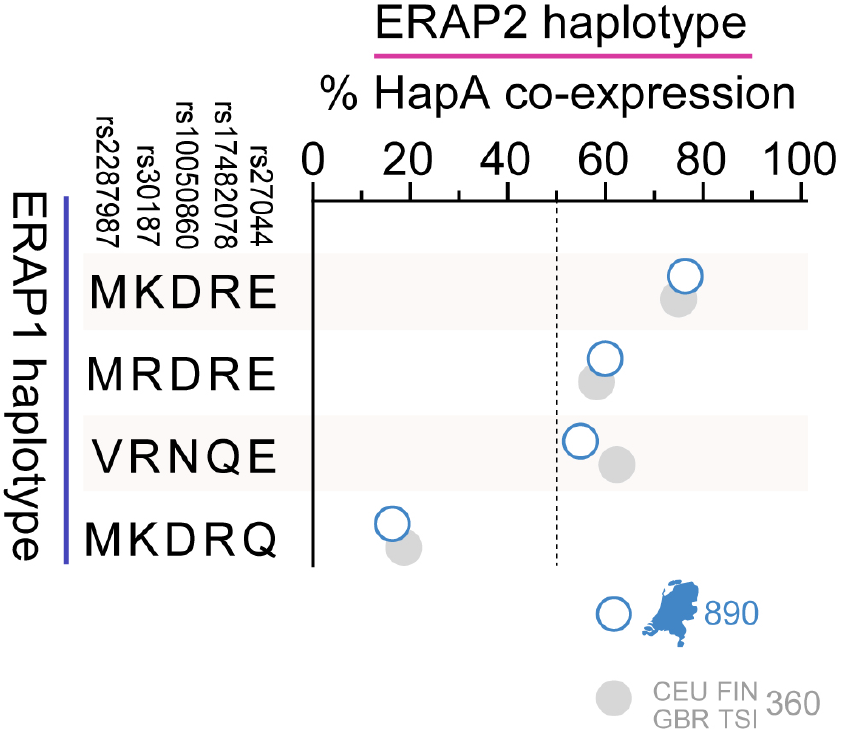
Non-random distribution of the ERAP2-protein coding haplotype across common ERAP1 haplotypes based on 5 SNPs. Five missense variants in *ERAP1* encode four discrete and common (>1% freq) haplotypes [12], while *ERAP2* encodes 1 common protein-coding haplotype (*HapA* tagged by the A allele of rs2248374) and non-coding haplotypes (tagged by the G allele). Each chromosome harbors 1 ERAP1 haplotype and 1 ERAP2 haplotype. We outlined the percentage of *HapA* per ERAP1 haplotype on the same chromosome using phased ERAP1-ERAP2 haplotype data of 890 Dutch controls (blue) and ERAP1-ERAP2 haplotype data for the 360 samples from the CEU, FIN, GBR, and TSI populations (grey) from the 1000 Genomes sample collection used for functional investigation in this study [48].

## Supplementary Tables

**Supplementary Table 1.** Association results for common functional variants at 5q15 in *ERAP1, ERAP2* and *LNPEP* genes in Dutch and Spanish populations.

| CHR | Gene | Function | SNP | Position | Allele<br>(minus<br>strand<br>) | Netherlands |  |  |  | Spain |  |  |  | Meta -analysis |  |
| --- | --- | --- | --- | --- | --- | --- | --- | --- | --- | --- | --- | --- | --- | --- | --- |
|  |  |  |  |  |  | Freq |  | OR (95% CI) | P | Freq |  | OR (95% CI) | P |  |  |
|  |  |  |  |  |  | Case<br>(n=84) | Control<br>(n=890) |  |  | Case<br>(n=46) | Control<br>(n=2103) |  |  | OR | P |
| 5 | ERAP1 | eQTL of ERAP1 | rs1057569 | 96109610 | A | 0.42 | 0.29 | 1.69(1.19-2.40) | 3.25 x 10<sup>-3</sup> | 0.35 | 0.28 | 1.37(0.89-2.11) | 0.15 | 1.56 | 1.41 x 10<sup>-3</sup> |
| 5 | ERAP1 | 730Q | rs27044 | 96118852 | G | 0.15 | 0.29 | 0.45(0.29-0.71) | 5.84 x 10<sup>-4</sup> | 0.24 | 0.32 | 0.67(0.42-1.08) | 0.10 | 0.55 | 2.89 x 10<sup>-4</sup> |
| 5 | ERAP1 | 725Q | rs17482078 | 96118866 | T | 0.36 | 0.19 | 2.20(1.51-3.20) | 3.53 x 10<sup>-5</sup> | 0.29 | 0.19 | 1.73(1.10-2.71) | 0.017 | 1.99 | 2.58 x 10<sup>-6</sup> |
| 5 | ERAP1 | 575N | rs10050860 | 96122210 | T | 0.38 | 0.21 | 2.24(1.56-3.24) | 1.51 x 10<sup>-5</sup> | 0.29 | 0.19 | 1.70(1.08-2.67) | 0.021 | 2.01 | 1.46 x 10<sup>-6</sup> |
| 5 | ERAP1 | 528K | rs30187 | 96124330 | T | 0.16 | 0.34 | 0.37(0.24-0.58) | 1.20 x 10<sup>-5</sup> | 0.35 | 0.40 | 0.80(0.52-1.23) | 0.31 | 0.55 | 1.53 x 10<sup>-4</sup> |
| 5 | ERAP1 | 349V | rs2287987 | 96129535 | C | 0.38 | 0.20 | 2.25(1.56-3.24) | 1.49 x 10<sup>-5</sup> | 0.29 | 0.19 | 1.70(1.09-2.67) | 0.020 | 2.01 | 1.41 x 10<sup>-6</sup> |
| 5 | ERAP1 | 346D | rs27895 | 96129543 | T | 0.07 | 0.07 | 0.92(0.47-1.81) | 0.82 | 0.05 | 0.06 | 0.86(0.35-2.11) | 0.75 | 0.90 | 0.70 |
| 5 | ERAP1 | 276M | rs26618 | 96130836 | C | 0.29 | 0.25 | 1.32(0.90-1.92) | 0.16 | 0.20 | 0.22 | 0.85(0.51-1.42) | 0.53 | 1.13 | 0.45 |
| 5 | ERAP1 | 127R | rs26653 | 96139250 | C | 0.17 | 0.28 | 0.50(0.33-0.78) | 1.87 x 10<sup>-3</sup> | 0.25 | 0.31 | 0.74(0.46-1.18) | 0.20 | 0.60 | 2.00 x 10<sup>-3</sup> |
| 5 | ERAP1 | 56K | rs3734016 | 96139464 | T | 0.05 | 0.04 | 1.42(0.68-2.95) | 0.35 | 0.04 | 0.03 | 1.69(0.62-4.58) | 0.31 | 1.51 | 0.18 |
| 5 | ERAP1 | 12I | rs72773968 | 96139595 | A | 0.06 | 0.13 | 0.48(0.24-0.94) | 0.03 | 0.14 | 0.15 | 0.90(0.50-1.61) | 0.72 | 0.69 | 0.09 |
| 5 | ERAP2 | Splice-variant | rs2248374 | 96235896 | A | 0.67 | 0.47 | 2.26(1.59-3.22) | 6.11 x 10<sup>-6</sup> | 0.56 | 0.48 | 1.40(0.92-2.12) | 0.12 | 1.85 | 7.96 x 10<sup>-6</sup> |
| 5 | LNPEP | eQTL of ERAP2 | rs10044354 | 96320495 | T | 0.63 | 0.42 | 2.42(1.69-3.45) | 1.21 x 10<sup>-6</sup> | 0.53 | 0.40 | 1.68(1.11-2.55) | 0.014 | 2.07 | 1.24 x 10<sup>-7</sup> |

**Supplementary Table 2.** Results of test for association of the haplotype rs2287987 (risk allele C) and rs10044354 (risk allele T) with Birdshot Uveitis in 130 cases and 2993 controls using a logistic regression model.

| Cohort | rs2287987 | rs10044354 | Haplotype Frequency |  | OR [95%CI] | P value |
| --- | --- | --- | --- | --- | --- | --- |
|  |  |  | Cases | Controls |  |  |
| Netherlands | C | T | 0.24 | 0.12 | 3.1 [1.96-4.90] | $1.4 \times 10^{-6}$ |
|  | T | T | 0.39 | 0.31 | 1.44 [0.99-2.08] | 0.05 |
|  | C | C | 0.14 | 0.08 | 1.34 [0.75-2.40] | 0.33 |
| | T | C | 0.23 | 0.49 | 0.30 [0.20-0.46] | $3.5 \times 10^{-8}$ |
| Spain | C | T | 0.19 | 0.11 | 2.29 [1.31-4.02] | $3.9 \times 10^{-3}$ |
|  | T | T | 0.34 | 0.30 | 1.23 [0.79-1.92] | 0.36 |
|  | C | C | 0.10 | 0.08 | 1.2 [0.57-2.53] | 0.63 |
| | T | C | 0.37 | 0.51 | 0.53 [0.34-0.84] | $6.2 \times 10^{-3}$ |
| Meta-analysis | C | T | | | 2.75 [1.92-3.92] | $2.6 \times 10^{-8}$ |
|  | T | T |  |  | 1.35 [1.02-1.80] | 0.04 |
|  | C | C |  |  | 1.291 [0.81-2.03] | 0.28 |
| | T | C | | | 0.39 [0.30-0.55] | $3.9 \times 10^{-9}$ |

**Supplementary Table 3.**
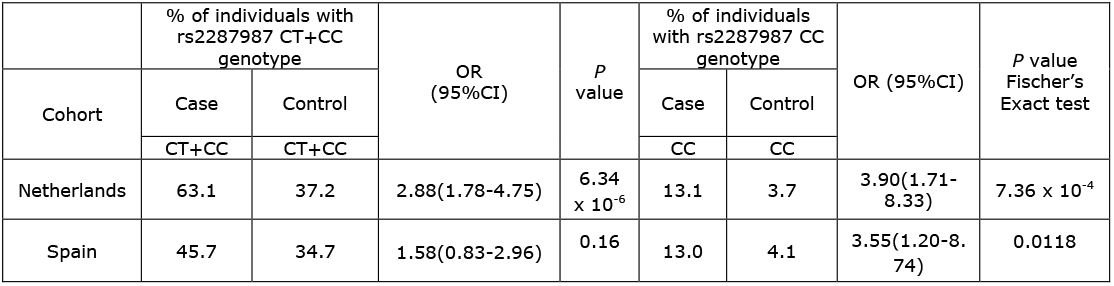
Distribution of Hap10 in the Dutch and Spanish cases and controls. The frequency of the rs2287987-C (i.e. Hap10+ individuals) or rs2287987-CC (i.e. Hap10 homozygous individuals) genotypes in cases and controls was tested by Fischer’s exact test. OR; odds ratio. 95%CI; 95% confidence interval.

**Supplementary Table 4.** Birdshot-associated SNPs are eQTLs for *ERAP1* and *ERAP2* isoforms. We tested the association between the risk alleles rs2287987 (correcting for rs1057569) and rs10044354 (correcting for rs2248374) with 4 detected transcripts at 5q15 in 360 lymphoid cell lines from European ancestry of the 1000 genomes project [48]. The incidence rate ratio (IRR) is calculated as the exponentiated genotype coefficient from the generalized linear mixed model and represents the change in transcript expression (scaled counts) associated with each sequential increase in risk allele count [23]. The IRR^2^ is conceptually similar to the change in expression in homozygous risk individuals relative to those homozygous for the protective allele (e.g. 1.542 ~ 54% increase) as previously described [23]. The mean count per transcript is indicated for individuals homozygous for the protective and risk allele. The *P* values are corrected for multiple testing using Bonferroni’s correction. Observations of IRR^2^ <0.60 or >1.30 (i.e. >30% change in expression) are considered biologically relevant and are highlighted in orange (increase) and blue (decrease).

|  | Risk allele | IRR (95%CI) | IRR<sup>2</sup> (95%CI) | % change expression AA x BB | P value | Mean count in homozygous Protective allele | Mean count in homozygous Protective allele | Ratio Risk/Protective AA x BB |
| --- | --- | --- | --- | --- | --- | --- | --- | --- |
| **ERAP1~201 ENST00000296754.7** |  |  |  |  |  |  |  |  |
| rs1057569 | A | 4.13(3.63 -4.7) | 17.05(13.15-22.1) | 1605% | 1.66 x 10<sup>-100</sup> | 1.17 | 15.78 | 13.49 |
| rs2287987 | C | 2.59(2.17 -3.09) | 6.7(4.7-9.56) | 570% | 1.56 x 10<sup>-24</sup> | 3.18 | 14.86 | 4.67 |
| rs2287987 corrected for rs1057569 | C | 1.03(0.87 -1.21) | 1.05(0.76-1.47) | 5% | 1 |  |  |  |
| rs10044354 corrected for rs1057569 | T | 1.01(0.9 -1.13) | 1.02(0.82-1.27) | 2% | 1 | 5.29 | 7.22 | 1.36 |
| **ERAP1~202 ENST00000443439.6** |  |  |  |  |  |  |  |  |
| rs1057569 | A | 0.62(0.59 -0.64) | 0.38(0.35-0.42) | -62% | 1.93 x 10<sup>-105</sup> | 74.37 | 25.36 | 0.34 |
| rs2287987 | C | 0.66(0.62 -0.7) | 0.44(0.39-0.49) | -56% | 1.77 x 10<sup>-44</sup> | 68.01 | 27.54 | 0.40 |
| rs2287987 corrected for rs1057569 | C | 0.98(0.91 -1.05) | 0.96(0.83-1.11) | -4% | 1 |  |  |  |
| rs10044354 corrected for rs1057569 | T | 0.93(0.9 -0.97) | 0.87(0.8-0.94) | -13% | 8.06 x 10<sup>-3</sup> | 64.10 | 49.61 | 0.77 |
| **ERAP2~202 ENST00000437043.7** |  |  |  |  |  |  |  |  |
| rs1057569 corrected for rs2248374 | A | 1.07(1 -1.14) | 1.14(1.01-1.3) | 14% | 0.54 | 57.05 | 69.31 | 1.21 |
| rs2287987 corrected for rs2248374 | C | 1.06(0.99 -1.14) | 1.13(0.99-1.29) | 13% | 1 | 57.05 | 72.10 | 1.26 |
| rs10044354 corrected for rs2248374 | T | 1.83(1.66 -2.02) | 3.35(2.76-4.06) | 235% | 3.48 x 10<sup>-33</sup> | 23.75 | 89.61 | 3.77 |
| **LNPEP~201 ENST00000231368.9** |  |  |  |  |  |  |  |  |
| rs1057569 | A | 0.99(0.95 -1.03) | 0.98(0.9-1.06) | -2% | 1 | 92.22 | 89.60 | 0.97 |
| rs2287987 | C | 0.99(0.94 -1.03) | 0.97(0.89-1.06) | -3% | 1 | 91.06 | 87.86 | 0.96 |
| rs10044354 | T | 0.97(0.93 -1.01) | 0.94(0.87-1.01) | -6% | 1 | 92.00 | 87.49 | 0.95 |

**Supplementary Table 5.** ERAP1 amino acids positions for Haplotype 10 (Hap10). Using molecular cloning, full-length *ERAP1* was sequenced from a patient homozygous for *Hap10 and a patient heterozygous for Hap10 (based on common genotyped SNPs)*. Previously reported amino acid positions in *ERAP1* are adapted from *Reeves et* a/.[24] and *Ombrello et al*. [21]

|  | Amino Acid position in ERAP1 |  |  |  |  |  |  |  |  |  |  |  |  |  |  |  |  |  |
| --- | --- | --- | --- | --- | --- | --- | --- | --- | --- | --- | --- | --- | --- | --- | --- | --- | --- | --- |
| Allotype | 12<sub>a</sub> | 56<sup>a</sup><sub>b</sub> | 82<sup>b</sup> | 102<sub>b</sub> | 115<sub>b</sub> | 127<sup>a</sup><sub>b</sub> | 199<sub>b</sub> | 276<sub>a,b</sub> | 346<sup>a,b</sup> | 349<sub>a,b</sub> | 528<sub>a,b</sub> | 575<sub>a,b</sub> | 725<sub>a,b</sub> | 727<sub>b</sub> | 730<sub>a,b</sub> | 737<sup>b</sup> | 752<sub>b</sub> | 847<sub>b</sub> |
| Hap10<sup>c</sup> | T | E | V | I | L | P | F | I | G | V | R | N | Q | L | E | V | R | M |
| *001<sup>d</sup> |  |  | V | I | L | P | F |  |  | V | R | N | Q | L | E | V | R | M |
| Hap10<sup>e</sup> |  | E |  |  |  | P |  | I | G | V | R | N | Q |  | E |  |  |  |

<sup>a</sup>Amino acid identified by genotyping/imputation
<sup>b</sup>Amino acid identified by sanger sequencing of full-length cDNA of *ERAP1*
<sup>c</sup> Full-length cDNA (Allotype) amino acid sequence obtained from two Birdshot uveitis patients heterozygous and homozygous for *Hap10* in this study.
<sup>d</sup>Allotype name and amino acid positions adapted from [24]
<sup>e</sup>Allotype name and amino acid positions adapted from [21]

**Supplementary Table 6.** The frequency of the rs2248374 A allele in *ERAP2* cooccurring with each of the 8 common ERAP1 haplotypes is non-random in phased haplotype data of 890 Dutch and 2103 Spanish controls. The *ERAP2* protein-coding haplotype [9] is tagged by the A allele of rs2248374 in *ERAP2*. We used a two-sided exact binominal test to assess if the observed frequencies were non-random (deviation from a test probability of 0.5 or the hypothesis that ERAP2 co-occurs in 50% of any particular ERAP1 haplotype). Frequency distribution of ERAP2 for each of the ERAP1 haplotypes was considered non-random if the frequency deviated at a significant level of [A or G rs2248374 for 8 ERAP1 haplotypes = 0.05/16] p<2.5 x 10^−3^ in both the Dutch and Spanish populations (with consistent direction of deviation from 50%). ERAP1 haplotypes that fulfilled these criteria are highlighted (in orange haplotypes that more frequently co-occur with rs2248374-A allele and in blue haplotypes that less frequently co-occur with rs2248374-A allele).

| ERAP1 haplotype | Number of Dutch haplotypes (n=1780) | % Co-occurring A-rs2248374 (95% CI) | P value | Number of Spanish haplotypes (n=4206) | % Co-occurring A-rs2248374 (95% CI) | P value |
| --- | --- | --- | --- | --- | --- | --- |
| *Hap1\** | 225 | 24(18-30) | $7.35 \times 10^{-16}$ | 641 | 43(39-47) | $3.71 \times 10^{-4}$ |
| *Hap2\** | 268 | 7(4-11) | $2.66 \times 10^{-52}$ | 683 | 34(30-37) | $1.02 \times 10^{-17}$ |
| *Hap3* | 101 | 76(67-84) | $1.18 \times 10^{-7}$ | 340 | 55(49-60) | 0.093 |
| *Hap5* | 117 | 30(22-39) | $1.65 \times 10^{-5}$ | 259 | 42(36-49) | 0.018 |
| *Hap6\** | 165 | 8(4-13) | $3.10 \times 10^{-31}$ | 348 | 35(30-40) | $1.42 \times 10^{-8}$ |
| *Hap7\** | 71 | 96(88-99) | $5.06 \times 10^{-17}$ | 110 | 72(62-80) | $5.34 \times 10^{-6}$ |
| *Hap8\** | 442 | 83(79-86) | $1.41 \times 10^{-46}$ | 935 | 61(58-64) | $5.33 \times 10^{-12}$ |
| *Hap10* | 345 | 55(49-60) | 0.085 | 808 | 51(47-54) | 0.65 |
| **Total (% of ERAP1 haplotypes)** | **1734 (97%)** |  |  | **4124 (98%)** |  |  |

